# Molecular Phylogenetics of HIV-1 Subtypes in African Populations: A Case Study of Sub-Saharan African Countries

**DOI:** 10.1101/2022.05.18.492401

**Authors:** Hesborn Omwandho Obura, Clement Dastan Mlay, Lindani Moyo, Brenda Muthoni Karumbo, Kauthar Mwanamkuu Omar, Erick Masunge Sinza, Gladys Jerono Rotich, Wilson Mudaki, Brenda Muthoni Kamau, Olaitan I. Awe, Allissa Dillman

## Abstract

Despite advances in antiretroviral therapy that have revolutionized HIV-1 disease management, effective control of the HIV-1 infection pandemic remains elusive. Increased HIV-1 infection rates and genetic diversity in Sub-Saharan African countries pose a challenge in HIV-1 clinical management. This study provides a picture of HIV-1 genetic diversity and its implications for HIV-1 disease spread and the effectiveness of therapies in Africa. Whole-genome sequences of HIV-1 were obtained from Genbank using the accession numbers from 10 African countries with high HIV-1 prevalence. Alignment composed of the query and reference sequences retrieved from the Los Alamos database. The alignment file was viewed and curated in Aliview. Phylogenetic analysis was done by constructing a phylogenetic tree using the maximum likelihood method implemented in IQ-TREE. The clustering pattern of the studied countries showed both homogeneous clustering and heterogeneous clustering with all Zambia sequences clustering with HIV-1 subtype-C indicating local distribution of only subtype-C. Sequences from nine countries showed heterogeneous clustering along with different subtypes as well as individual clustering of the sequences away from references suggesting cross border genetic exchange. Sequences from Kenya and Nigeria clustered with almost all the HIV-1 subtypes suggesting high HIV-1 genetic diversity in Kenya and Nigeria as compared to other African countries. Our results indicate that there is the presence of subtype-specific HIV-1 polymorphisms and interactions during border movements.

## 1 INTRODUCTION

Human immunodeficiency virus (HIV) and many simian immunodeficiency viruses (SIV) that are prevalent in Africa and affect the continent’s primate species belong to the family of *Retroviridae* and genus *Lentivirus* (Daw *et al*., 2017). HIV is categorized into HIV type 1 (HIV-1) and HIV type 2 (HIV-2) based on differences in the viral antigens and genetic characteristics (Seitz, 2016). HIV displays important genetic variability with each HIV-1 replication cycle generating approximately one mutation per genome (Giovanetti *et al*., 2020). The differences between HIV-1 and HIV-2 are fairly well documented in terms of transmissibility, pathogenesis and pattern of spread (Santoro *et al*., 2013).

HIV-1 is believed to have originated in non-human primates in West-central Africa through a process known as zoonosis, and it transferred to humans in the early 20th century (Giovanetti *et al*., 2020). Retrospective studies trace the origin of HIV-1 to the Democratic Republic of Congo(Hayflick, 1992) and is subsequently spreading to other areas in sub-Saharan Africa and West Africa. HIV-1 diversified extensively during worldwide dissemination, undergoing a constant molecular evolution. HIV-1 is more virulent and the major cause for the global AIDS pandemic (MA *et al*., 2017) while HIV-2 is less virulent (Kanki *et al*., 1994) and less common. HIV-1 is categorized into four groups (M, N, O and P) (Kotaki *et al*., 2015), each of which is a derivative of simian immunodeficiency viruses that naturally infect chimpanzees (SIVcpz) (Sharp and Hahn, 2011). The HIV-1 group M (Major) alone is responsible for more than 95% of the AIDS pandemic (Simon *et al*., 1998) and has been used as a representative of most studies conducted (Adhiambo *et al*., 2021).

Genetic variation in HIV-1 is caused by error-prone reverse transcriptase enzymes, recombination events during replication of the virus, HIV-1 rapid turnover in the body and immune system selective pressures (Désiré *et al*., 2018). The molecular basis of HIV-1 variability is the highly error-prone reverse transcriptase enzyme (Roberts *et al*., 1988) while its recombination is a mechanism used by the virus for survival to escape the body’s defence mechanism (Burke, 1997). The rate of nucleotide substitutions introduced by the reverse transcriptase is approximately 10^−4^ per nucleotide per cycle of replication, equivalent to one nucleotide substitution per genome during a single replication cycle (Bridgette and Connell, 2007). Insertions, deletions and duplications also contribute to the genetic heterogeneity of HIV-1(Bernstein *et al*., 2018).

HIV-1 has a rapid turnover with approximately 10^9^ virions per day generated in an HIV-1 infected individual. The half-life of plasma virus producing cell is estimated to two and an almost complete replacement of wild-type strains by drug resistant virus occurs in plasma within 2–4weeks (Santoro and Perno, 2013). The emergence of HIV-1 variants with antiretroviral resistance is attributed to the rapid viral turnover in combination with a high mutation rate during antiretroviral treatment (Varmus, 1987). This process contributes strongly to high level multiple drug resistance (Moutouh *et al*., 1996). Each retroviral particle contains two copies of single-stranded RNA, and template switches occur frequently during reverse transcription, generating mutations and recombination by intramolecular and intermolecular jumps (Onafuwa *et al*., 2003).

Recombination triggers drug resistant mutations in HIV-1, leading to increased resistance to a particular drug (Moradigaravand *et al*., 2014), or the generation of multidrug resistant variants (Moutouh *et al*., 1996). In addition, recombination may lead to mutations that compensate for a loss in viral fitness or replicative capacity due to previous acquisition of resistance mutations (Agudelo-Rojas *et al*., 2019; Moradigaravand *et al*., 2014; Moutouh *et al*., 1996; Song *et al*., 2018). This interaction between recombination, mutations, and viral fitness is highly intricate, but nonetheless, recombination and its mechanisms, especially at the level of diverse subtypes, warrant further investigation. The potential for genetic differences among subtypes to yield different patterns of resistance-conferring mutations is supported by natural variation among HIV subtypes in genetic content (40% variation in the env gene, and 8–10% variation in the pol/gag genes) (Santoro and Perno, 2013). This issue acquires special relevance in view of the fact that the HIV pol gene is the major target for all major classes of anti-HIV drugs and most HIV strains show hotspots for recombination in *gag-pol* and *env* regions.

HIV-1 genetic variability is the major obstacle in the treatment of HIV and development of effective drugs (Santoro & Perno, 2013). The increased genetic variation of HIV-1 presents clinical and public health challenges (Sides *et al*., 2005). Precisely, patients monitoring, treatment, diagnostic testing, epidemiologic surveillance and drug development are influenced by HIV-1 genetic diversity (Sides *et al*., 2005). HIV-1 infections have been managed with highly active antiretroviral therapy; the nucleoside and non-nucleoside reverse transcriptase inhibitors and protease inhibitors (Kwara *et al*., 2005), with countries like Kenya scaling up antiretroviral therapy (ART) (Borgdorff *et al*., 2018; Ayah, 2018). However, with increased uptake of ART, there would be drug resistant strains among ART exposed individuals and there will be subsequent increase in drug resistance mutations (Wallis *et al*., 2014).

High HIV-1 infection rates and genetic diversity especially in the African population pose significant challenges in HIV-1 clinical management and drug design, as well as training the healthcare system and the economy. The new strains identified may confer resistance to antiretroviral drugs making it difficult to manage the HIV-1 epidemic. Current and extensive data on HIV-1 genetic diversity across Sub-Saharan countries is needed. In this study, we want to identify the various HIV-1 subtypes circulating in Africa to give a recent characterization of the HIV-1 epidemic in African populations which will be integral in informing health policy. The study findings will also provide useful insights to the natural history of HIV at the molecular, host and community levels, which is a critical focal point for the assessment of the effectiveness of treatment, intervention strategies and informing public health policy.

This study is therefore aimed at identifying the various HIV-1 subtypes circulating in the Sub-Saharan Africa region and to provide information on the genetic diversity of HIV-1 in this region. These findings will also provide useful insights critical for the assessment of the effectiveness of treatment, intervention strategies and informing public health policy.

The proposed research is aimed at *in silico* screening of HIV-1 whole genome sequences from NCBI virus repository from African countries and genetic characterization using computational approaches.

The objective is to investigate the prevalence of viral variants circulating in Sub-Saharan African countries to better characterize the HIV-1 epidemic in African populations.

## 2 METHODS

### 2.1 Workflow

### 2.2 Sequence Acquisition and Retrieval

Whole genome sequences of HIV-1 isolates from infected patients in 10 selected African countries were retrieved from GenBank using a custom python script (appendix I). The selected African countries are: Kenya, South Africa, Tanzania, Nigeria, Cameroon, Guinea Bissau, Uganda, Rwanda, Ethiopia and Zambia. The selected African countries are highly populated and have a high prevalence of HIV infection according to WHO report (https://www.afro.who.int/health-topics/hivaids). A total of 150 sequences were analysed, representing 15 sequences from each selected African country. In addition, 16 reference sequences of various HIV-1 subtypes were included in the analysis as reported previously (Adhiambo *et al*., 2021).

### 2.3 Sequence Alignment

Alignment composed of the query and reference sequences, were constructed using MAFFT by specifying the input file, the output file and the output file format described by (Katoh & Standley, 2013).

### 2.4 Curation

The alignment file was visualised and curated in AliView. The gaps that were shared by most sequences (more than 50% of the sequences) were removed; all the positions with gaps and any adjacent unambiguously aligned positions were deleted. The terminal ends were also trimmed to have similar length for sequences used in the phylogenetic analysis.

### 2.5 Phylogenetic Analysis

The taxonomic clusters were constructed and identified using the maximum-likelihood approach implemented by the IQ-TREE version 2.1.4 with 1000 replicates for bootstrap and the statistical significance of the phylogenetic tree topology was estimated using bootstrap analysis. The best substitution model, GTR+F+R9, was chosen based on BIC and AICcs. The phylogenetic tree was then visualized and manipulated using Figtree version 1.44.

### 2.6 Implementation

The workflow used in the study was implemented using the following tools:

#### 2.6.1 MAFFT

Command line-based MAFFT was used for alignment of the sequences to the reference genome. The input format for this tool is Fasta format. The type of input sequences (amino acid or nucleotide) is automatically recognized. The output format used was fasta while the strategy used was auto. The MAFFT command was run with no additional arguments (Katoh *et al*., 2018).

MAFFT is open-source software and installation details can be found at this linke: https://mafft.cbrc.jp/alignment/software/. It can also be installed via the terminal using the conda environment by simply running the following commands:

~~~
      conda install -c bioconda mafft
      or
      conda install -c bioconda/label/cf201901 mafft
      Usage: **mafft [arguments] input > output**
~~~

#### 2.6.2 ALIVIEW

AliView is an alignment viewer and editor designed to meet the requirements of Next-Generation Sequencing era phylogenetic datasets. AliView handles alignments of unlimited size in the formats most commonly used, i.e. FASTA, Phylip, Nexus, Clustal and MSF. The intuitive graphical interface makes it easy to inspect, sort, delete, merge and realign sequences as part of the manual filtering process of large datasets. AliView also works as an easy-to-use alignment editor for small as well as large datasets (Larsson,2014).

AliView is released as open-source software under the GNU General Public License, version 3.0 (GPLv3), and is available at GitHub (www.github.com/AliView).

The program is cross-platform and extensively tested on Linux, Mac OS and Windows systems. Downloads and help are available at http://ormbunkar.se/aliview

#### 2.6.3 IQ-TREE

IQ-TREE takes as input multiple sequence alignment and will reconstruct an evolutionary tree that is best explained by the input data from MAFFT.

IQ-TREE is available at http://www.iqtree.org/ Development: https://github.com/iqtree/iqtree2

It can also be installed using conda by running the following commands in the terminal:

~~~
     conda install -c bioconda iqtree
     or
     conda install -c bioconda/label/cf201901 iqtree
     Usage: **iqtree -s [MSA file]**
~~~

#### 2.6.4 FigTree

FIGTREE takes as input the tree file and shows a visualization of the phylogenetic tree (FigTree, 2022). The phylogenetic tree was rooted via midpoint rooting and presented in a radial layout.

Compiled binaries (for Mac, Windows and Linux) are available here: https://github.com/rambaut/figtree/releases.

Source code is available on GitHub.

It can also be installed using conda by running the following commands in the terminal:

~~~
 conda install -c bioconda figtree
     Usage: **figtree [tree file]**
~~~

## 3 RESULTS

Phylogenetic results in Figure 2 above consists of 150 sequences from 10 different countries in Africa and reference sequences of each HIV-1 subtype (A1,A2,B,C,D,G, AB, and AD) retrieved from Los Alamos Database. The clustering pattern of the studied countries showed both homogeneous and heterogeneous clustering of HIV-1. All sequences from Zambia (100%), thirteen sequences from South Africa (86.67%) indicated local diversity, twelve sequences from Ethiopia (80%), four sequences from Tanzania (26.67%) and two sequences from Kenya (13.33%) clustered together with subtype-C. Five sequences from Uganda (33.33%), and three sequences from Tanzania (20%) clustered together with subtype-D. Two sequences from South Africa (13.22%) and one sequence from Guinea Bissau (6.67%) clustered together with subtype-B. Ten sequences from Kenya (66.67%), five sequences from Tanzania(33.3%), one sequence from Nigeria (6.67%) and seven sequences from Rwanda (46.67%) clustered together with subtype-A1. Two sequences from Uganda (13.33) were identified as recombinants by clustering together with subtype-AD. Three sequences from Nigeria (20%) and two sequences from Cameroon (13.22%) clustered together with subtype-A2. Seven sequences from Nigeria (46.67%) and two sequences from Cameroon (13.22%) clustered together with subtype-G (Fig. 2). Two sequences from Guinea Bissau (13.22%) and one sequence from Ethiopia (6.67%) clustered together and were close to subtype-G. Six sequences from Cameroon (40%) did not cluster with all the HIV-1 subtype. Three sequences from Nigeria (20%), two sequences from Cameroon (13.22%) and two sequences from Guinea Bissau (13.22%) clustered in the same clade and close to the six sequences from Cameroon. Four sequences from Guinea Bissau (26.67%) clustered away from all other sequences from all countries and formed their own cluster (Fig. 2).

**Fig 1.**
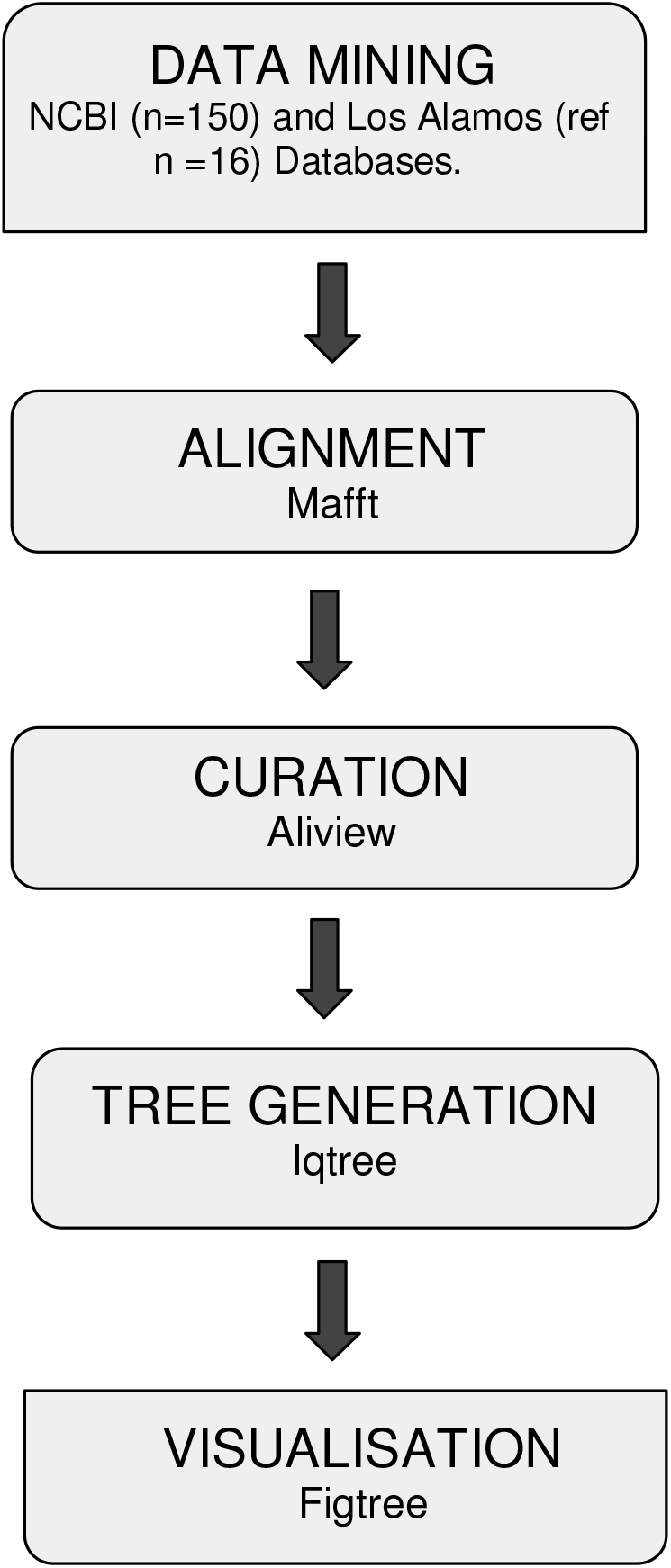
A summary of the computational workflow for the phylogenetic characterization of the human immunodeficiency virus (HIV) in Africa.

**Fig 2.**
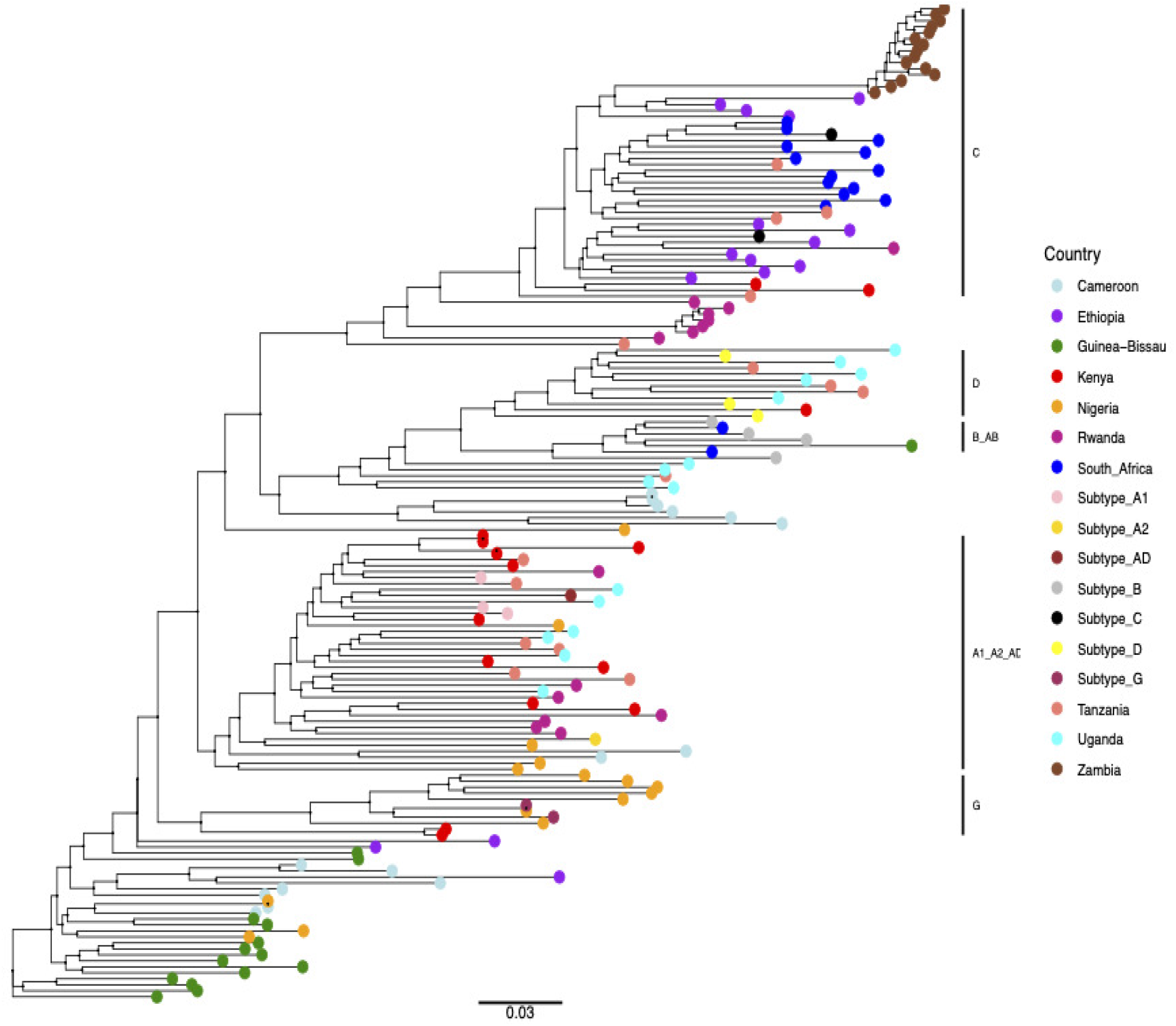
A phylogenetic tree showing the clustering of HIV-1 strain isolates in Sub-Saharan African countries and HIV-1 subtypes based on whole-genome sequences. Colours indicate each subtype and specific countries while vertical lines on the figure represent HIV-1 subtype clustering.

For the 150 sequences used for phylogenetic analysis, 128 (85.33%) showed close clustering with the HIV-1 subtypes references (A1, A2, C, D, G) and their recombinants (AD, AB) retrieved from Los Alamos HIV database. There were 28 sequences (14.67%) that clustered away from HIV-1 references but were phylogenetically closely related to sequences from different countries. This result is attributed to the limitation of this study in which not all of the HIV-1 subtype references were used. In this study subtype F, H, J and K were missing.

These similarities suggest an evolutionary trend and virus transmission within the population across the countries with an exception of Zambian sequences. This shows that cross border movement attributes to the introduction of new viral strains as observed in Kenya and Nigeria. The results of this study show that whole genome sequences have intrinsic genetic variation for phylogenetic analysis by the reconstruction of transmission histories compared to partial genes (Kaye *et al*., 2008).

## 4 DISCUSSION

An increased diversity or circulation of HIV-1 subtypes has been reported in this study as compared to previous studies across African countries and this poses a threat. This was because the emergence of certain subtypes or recombinant strains has differing outcomes on rates of transmission, disease progression and development of drug resistance. The new recombinants indicate that there could be new subtypes being introduced and circulating among different African countries due to cross-border movement of people from different regions of Africa and beyond. It is reassuring that current antiretroviral strategies appear to be effective against a broad spectrum of HIV-1 subtypes. However, further studies in larger and homogenous populations infected by different subtypes are required to better evaluate the viral diversity and to further elucidate viral polymorphisms and properties associated with transmission and disease progression may lead to new approaches to disease prevention and treatment. Efforts should be undertaken in diverse human populations infected with a variety of HIV-1 subtypes in order to fully understand the complex interplay of HIV and host in AIDS pathogenesis.

## ACKNOWLEDGEMENTS

The authors are indebted to the omics-focused codeathon organized by the African Society for Bioinformatics and Computational Biology (ASBCB) with support from National Institutes of health (NIH) office of data science strategy (ODSS): Oct 7 – 10, 2021.The authors thank Brigit Shea Sullivan, NIH Library Editing Service, for manuscript editing assistance. The authors also acknowledge Evans Mudibo for his immense support during this work.

## Author Contributions

*Conceptualization:* Hesborn Obura;

*Data Curation:* Brenda Muthoni, Clement Mlay, Hesborn Obura, Lindani Moyo, Gladys Rotich *Formal Analysis:* Clement Mlay, Brenda Muthoni, Hesborn Obura, Lindani Moyo, Kauthar Omar, Gladys Rotich

*Funding Acquisition:* Olaitan I. Awe, Allissa Dillman

*Investigation:* Hesborn Obura

*Methodology:* Hesborn Obura, Brenda Muthoni, Clement Mlay, Lindani Moyo, Gladys Rotich

*Project Administration:* Hesborn Obura, Lindani Moyo

*Resources:* Clement Mlay, Brenda Muthoni, Hesborn Obura, Lindani Moyo, Kauthar Omar Wilson mudaki, Gladys Rotich, Olaitan I. Awe

*Software:* Clement Mlay, Brenda Muthoni, Hesborn Obura *Supervision:* Hesborn Obura, Lindani Moyo, Olaitan I. Awe *Validation:* Hesborn Obura, Lindani Moyo

*Visualization:* Hesborn Obura, Lindani Moyo

*Writing – Original Draft Preparation:* Hesborn Obura, Lindani Moyo, Kauthar Omar *Writing – Review & Editing:* Lindani Moyo, Kauthar Omar, Hesborn Obura, Clement Mlay, Olaitan I. Awe

## Appendix I Sequence Search Script

The script searches for sequences from NCBI databases https://github.com/omicscodeathon/HIV-1-Genetics-Africans/blob/main/workflow/scripts/sequence_search.py

## Appendix II Sequence Download Script

The script downloads sequences from NCBI databases. https://github.com/omicscodeathon/HIV-1-Genetics-Africans/blob/main/workflow/scripts/data_downloader.py

## Appendix III Fasta File Edit Script

The script filters out the definition line variables to only select the accession number and country.

https://github.com/omicscodeathon/HIV-1-Genetics-Africans/blob/main/workflow/scripts/fasta_edit.py

## Appendix IV Phylogenetic Tree Generation Script

The script aligns sequences using the mafft tool, curates and inspects the sequences using aliview tool. It generates a phylogenetic tree using iqtree tool and views the tree using figtree tool. https://github.com/omicscodeathon/HIV-1-Genetics-Africans/blob/main/workflow/scripts/phylogenetic.sh

